# Straightforward and reproducible analysis of bacterial pangenomes using Pagoo

**DOI:** 10.1101/2020.07.29.226951

**Authors:** Ignacio Ferrés, Gregorio Iraola

**Affiliations:** Microbial Genomics Laboratory, Institut Pasteur Montevideo, Montevideo, Uruguay; Center for Integrative Biology, Universidad Mayor, Santiago de Chile, Chile; Wellcome Sanger Institute, Hinxton, United Kingdom

## Abstract

Pangenome analysis is fundamental to explore evolutionary processes occurring in bacterial populations. However, the lack of standardized methods for handling diverse pangenomic datasets and complex metadata hinders more straightforward and reproducible downstream analyses. To fill this gap, we introduce Pagoo, a new framework that integrates pangenome data, analytical methods and visualization tools in a single object that can be easily stored, shared and responsively queried for improved biological interpretation of bacterial evolution.

---

The exponentially growing number of diverse bacterial genomes has prompted pangenome reconstruction as a gold standard to explore genetic diversity of bacterial populations^1,2^. Pangenome comparisons reveal genome evolutionary dynamics associated with important biological processes such as speciation, host-adaptation, pathogenicity or the acquisition of antimicrobial resistance. Pangenome reconstruction is typically performed from genes annotated in a set of whole-genome sequences. In general, coding sequences of different strains are grouped in orthologous clusters based on different similarity criteria. Then, pangenome data informs about the belonging of each gene encoded in each genome to a certain orthologous cluster. In the recent years, several software tools have been developed to reconstruct bacterial pangenomes, such as Roary^3^, panX^4^ or PanOCT^5^. These tools focus on automation of steps and optimization of computational costs to cluster thousands of sequences of increasingly large genomic datasets. However, there is a lack of tools that can take the output of pangenome reconstruction softwares and provide standardized and straightforward methods for data integration, storage, analysis and visualization.

Here, we introduce Pagoo, the first pangenome post-processing tool that can take the output of pangenome reconstruction softwares providing a standardized framework for its analysis. Pagoo is based on an object-oriented design built on a novel class system in R which implements: i) an integrative data structure for standardized storage of pangenome information such as orthologous clusters, sequences, annotations and metadata in a single object; ii) a set of straightforward methods for responsive querying, handling and subsetting of this data structure; and iii) a set of standard statistics and active visualizations leveraging flexible downstream comparative analyses. Along with extensive documentation, we show how Pagoo interacts with other widely used microbial genomics tools and the R ecosystem for improved analysis of bacterial populations.

A pangenome can be represented as individual genes which belongs to organisms (genomes) and that are also assigned to a cluster of orthologous genes. Pagoo stores this as a three-column matrix, with one column identifying an individual gene, the next one identifying the organism that this gene belongs to, and the last one identifying the orthologous cluster that the gene was assigned by the pangenome reconstruction method. Optionally, this matrix can contain additional columns as gene-specific metadata like annotations or functional assignments. Orthologous clusters and organisms can also take metadata represented as two different matrices, with the condition that each one must contain a column that correctly maps each observation (cluster or organism) into the former matrix. Gene sequences can also be added to this structure, with the condition that their names must also map to rows in the first matrix (Fig. 1A). This relational structure optimizes data storage avoiding duplication, enables flexibility for working with different data types and facilitates complex querying and analysis.

**Figure 1.**
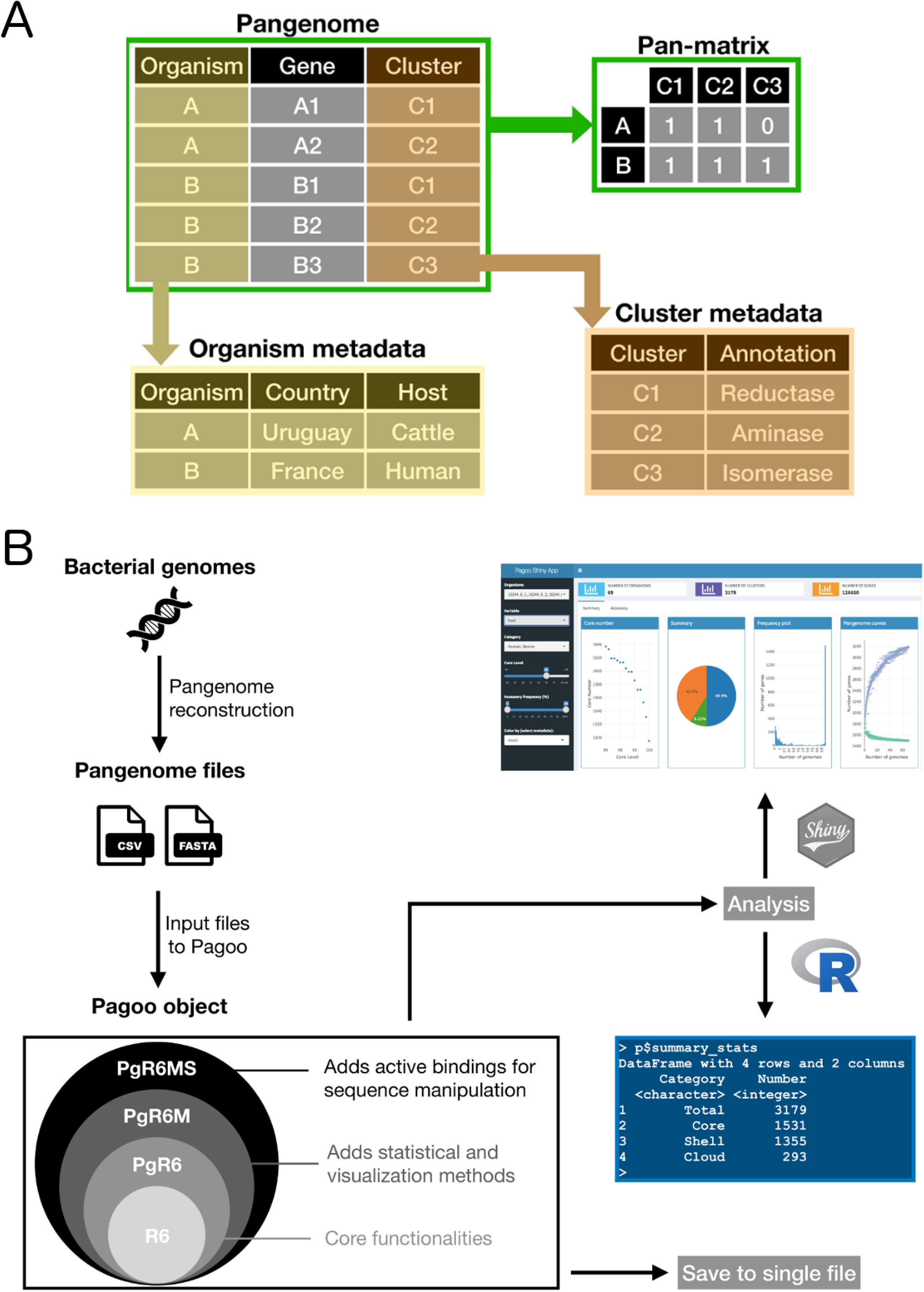
Framework and overall design of Pagoo. A) Example of the relational structure implemented to store, link and operate over different pangenome data types. B) General description of the workflow from assembled genomes to Pagoo analysis. Once pangenome files are created with any available pangenome reconstruction software, these files can be loaded to create the Pagoo object. The specific R6 classes store and manage different data types that allow to store all the information in a single file or perform comparative analyses using the R console interface or the Pagoo Shiny application.

Indeed, a salient and unique feature of Pagoo is that this data structure is stored and managed in an encapsulated, object-oriented fashion using the R6 package as backend. In contrast with traditional R programming, the R6 paradigm considers that methods belong to objects rather than to generic functions, so an object contains both the data and embedded methods to analyze it. In this context, the Pagoo object is built on three novel R6 classes. PgR6 is the most basic class that contains methods and functions for data handling and subsetting. Then, PgR6M inherits all the methods and fields from PgR6 and incorporates statistical methods and visualization tools based on the ggplot2 package^6^. PgR6MS inherits all capabilities from the others and adds methods for manipulation of DNA sequences using the Biostrings package^7^ (Fig. 1B). These classes support the main data types that typically represent a pangenome, providing a novel and synergistic framework to manage both the raw data and methods to perform operations and explore results with customized visualizations. Moreover, any of these classes could be further inherited and easily extended by third party applications.

Another remarkable feature of Pagoo is that raw data stored in the pangenome object is kept unaltered in the background, while users can query, mutate or subset the object using active bindings. This allows changing the state of the object without altering the original data. For example, users can temporarily hide certain organisms from the dataset, actively set thresholds that change the definition of core genes, or extract specific information from organisms, genes, clusters or sequences. Class-specific methods for generic subset operators are also implemented enabling seemly extraction of relevant field subsets straight from the object by using widely known R subset notation. Also, Pagoo provides specific methods to automatically generate the pangenome object from output files produced by standard pangenome reconstruction tools like Roary, and to save any changes to the object along with the unaltered original data as a single file. Importantly, Pagoo lacks of external dependencies and is built and tested in all three major operative systems (Linux, Windows and Mac). A detailed explanation of each method and operator for data input, saving and loading the pangenome object, and for specific data handling and subsetting is provided in the online user manual (https://iferres.github.io/pagoo/). Together, this implementation represents a new concept for pangenome data handling, facilitating reproducibility and enabling multiple and flexible analyses.

Pagoo also includes statistical and visualization methods. Customized plots and statistical analyses can be generated directly from the pangenome object using active bindings on the console or by deploying a built-in R-Shiny application. This interactive application is divided in two main components: (i) a general dashboard that interactively displays summary statistics including number of organisms, orthologous clusters and genes, core and accessory genome sizes, gene frequency barplots, pangenome curves and scrollable information about core genome clusters and genes (i.e. annotation or any other metadata); and (ii) a specific dashboard showing clustering of genomes according to accessory gene distances and Principal Components Analysis, genome-specific accessory genome sizes, visualization of gene presence/absence matrix with associated metadata and information about accessory gene clusters (Supplementary Information; Fig. S1). This interactive application allows responsive exploration of evolutionary trends in bacterial populations to guide downstream analyses, leveraging the interaction of Pagoo with other tools.

Remarkably, more complex comparative pangenome analyses can be performed by applying concise code recipes. We define recipes as relatively short snippets that pipe pangenome information extracted from the object as input to other R tools. We have developed example recipes (available in the online user manual at https://iferres.github.io/pagoo/articles/6-Recipes.html) to build core genome phylogenies, identify population structure, explore genome-wide selective pressures acting over the core genes and compare individual gene sequences against specific databases. Importantly, the development and implementation of recipes enable full reproducibility of publication-quality figures generated directly from the pangenome object (Fig. 2).

**Figure 2.**
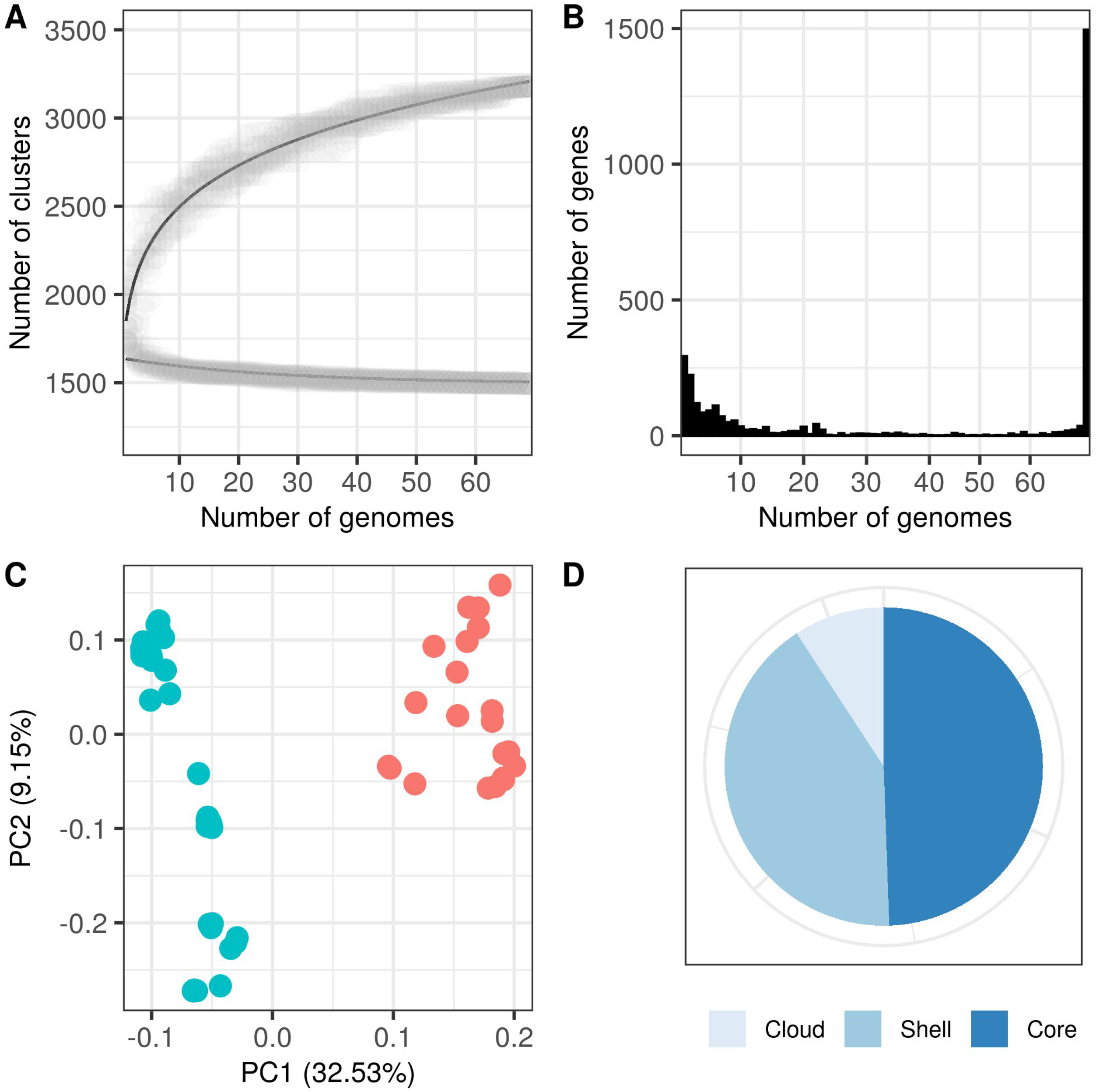
Results extracted from the pangenome object. Exploration of the *C. fetus* pangenome using information directly extracted from the pangenome object and customized aesthetics. Panel (A) shows pangenome and core genome curves with grey circles representing different sub-samples at increasing number of genomes; the black lines show the fitting to the power law and exponential decay functions, respectively. Panel B shows the distribution of genes in different subset of genomes. Panel C shows a Principal Components Analysis generated from the gene presence/absence matrix that clearly two groups of genomes, representing human-derived strains (red) and bovine-derived strains (green). Panel D shows the distribution of the pangenome in core genes and accessory genes (shell and cloud genes).

As a working example we used Pagoo to reanalyze a previously published study on the evolution of *Campylobacter fetus* pangenome. This species has a strong population structure with different lineages adapted to livestock or humans^8^. Briefly, we reconstructed a pangenome from 69 selected *C. fetus* genomes with Roary^3^ using default parameters and used its output to build a Pagoo object. Then, we performed a comparative analysis between livestock- and human-derived *C. fetus* genomes. The dynamic exploration of results using the Pagoo Shiny application (https://microgenlab.shinyapps.io/pagoo_campylobacter/) allowed us to recover main diversity patterns reported for this species, such as a marked difference between accessory genome size and gene presence/absence patterns between livestock- and human-adapted strains.

The advent of high-throughput sequencing technologies more than fifteen years ago pushed microbiology towards the field of comparative genomics, that rapidly transitioned from studies including few to thousands of genomes^2^. This substantially increased the complexity of datasets, requiring new approaches to systematically handle and track different components of interrelated pangenomic data. Pagoo introduces a new framework underpinned in a concept that leverages the simplicity of storing all the information in a standardized and reproducible manner in a single, shareable object. Along with future developments and addons, Pagoo aims to improve and facilitate current practices on the genomic analysis of bacterial populations.

## Acknowledgments

We thank Pablo Fresia and Daniela Costa for insightful comments and suggestions during testing of Pagoo. I.F. is funded by grant ANII-POS_NAC_2018_1_151494 from Agencia Nacional de Investigación e Innovación (ANII), Uruguay.

## Author contributions

G.I. and I.F. conceived the idea, I.F. developed the software and performed experiments, G.I. and I.F. wrote the manuscript.

