## Supplementary Information; Fig. S1 for "Straightforward and reproducible analysis of bacterial pangenomes using Pagoo"

A)

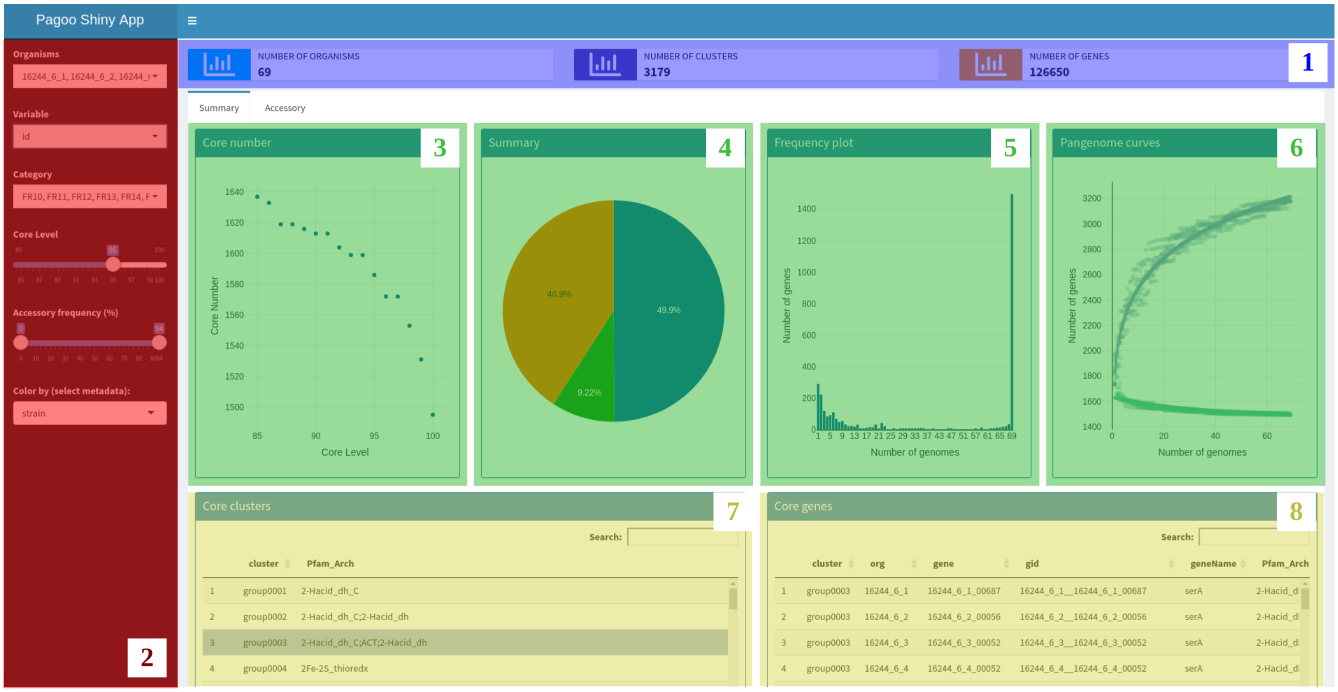


B)

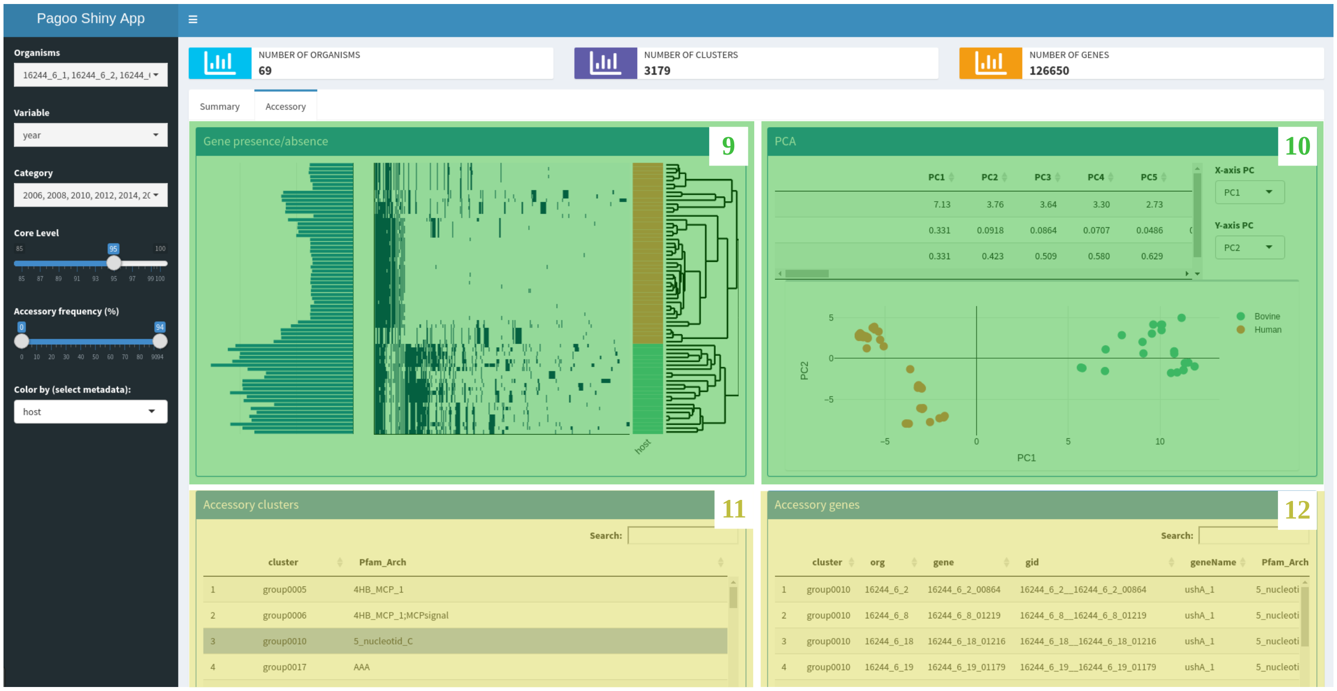


**Supplementary Figure 1. Overview of Pagoo Shiny Application.** A) Dashboard showing summary statistics and core genome characteristics of the dataset under analysis. B) Dashboard showing summary statistics and accessory genome characteristics of the dataset under analysis. A detailed description of each numbered panel is provided in the Supplementary Note.

**Supplementary Note**

**Detailed explanation of the Pagoo Shiny Application dashboards.** The Pagoo Shiny Application is composed by two main dashboards and one menu to modify parameters that allow responsive exploration of pangenome information. Here we explain the information displayed in each panel numbered in the Supplementary Figure 1:

<u>Panel A:</u> “Summary” tab

1) This header is permanently displayed and shows the number of organisms the number of orthologous clusters and the number of individual genes under analysis.

2) The left menu has several dropdown and scrollable menus that control parameters whose values affect plots and information displayed in the dashboard. The “Organisms” menu lists names for all organisms in the analysis and the user can select or deselect them. If the user deselects one ore several organisms, this will be excluded from the analysis and plots will be automatically updated without that information. The “Variable” menu lists all metadata columns associated to organisms present in the pangenome object. This menu is associated with the “Category” menu which displays all possible values for any selected variable. For example, in the provided dataset, the user can select “host” in the “Variable” menu so the “Category” menu will display the host of each organism. Then, the user can filter organisms for example by keeping only those from human host by deselecting the “Bovine” value. The “Core level” slide bar controls the percentage of organisms that must contain a certain gene for this to be considered a core genes. For example, if the core level is set to 90, a certain gene needs to be present in at least 90% of organisms (in this case 62 out of 69) to be counted as a core gene. The “Accessory frequency” slide bar and the “Color by” menu are explained in the “Panel B” section.

3) The “Core number” panel displays the number of retained core genes at different core levels. As expected, a more stringent core level (near 100%) will result in a smaller set of core genes. Hovering over dots will display a label in the format “(95, 1586)”, with the first number being the core level and the second being the number of core genes.

4) The “Summary” pie chart shows the percentage of core genes defined as those who appear in a frequency higher than the core level, the percentage of shell genes defined as those accessory genes present in more than 1 genome, and the percentage of cloud genes defined as genome specific genes. Hovering over the pie char will display a label showing the number of genes and percentages corresponding to these subsets.

5) The “Frequency plot” displays the number of genes while adding genomes in the dataset. This plot typically shows a “U” shape that describes the distribution of genes in the pangenome.

6) The “Pangenome curves” plot shows the cumulative number of genes as genomes are added in the dataset that tend to adjust to a power law function. This is useful to rapidly explore if a certain pangenome is open or close. This plot also shows the number of core genes as genomes are added in the dataset, that tend to adjust to an exponential decay function.

7) The “Core clusters” scrollable menu lists names of all core genome clusters and displays associated metadata like cluster annotation, etc.

8) The “Core genes” scrollable menu responds to the previous “Core clusters” menu. Once a certain core cluster is selected, those individual core genes belonging to the cluster are

displayed in the “Core genes” menu. For each gene, information like start and end position in the genome, annotation, gene name and strand are displayed.

Panel B: “Accessory” tab

In the left menu, the “Accessory frequency” slide bar controls the minimum and maximum frequency for any certain accessory gene to be considered in the analyses described as follows.

9) The “Gene presence/absence” plot displays the gene presence/absence matrix highlighting present accessory genes in each genome in blue and absent genes in white. The rows (organisms) are ordered according to a clustering analysis displayed in the right side that is based on the Bray-Curtis distance calculated over the presence/absence matrix. The bar plots in the left shows the number of the accessory genes in each genome. A vertical colored strip highlights each genome according to any metadata that the user selects in the “Color by” menu. Gene presence/absence blocks can be explored in detail by selecting a desired area of the plot that will zoom in automatically.

10) The “PCA” plot is based on the sample gene presence/absence matrix but user can select the principal components to be displayed. Also, information about the contribution of each principal component is showed in the top. Points can be colored by selecting metadata in the “Color by menu”.

11) “Accessory clusters” and 12) “Accessory genes” works as explained for 7 and 8, respectively.
